# The apportionment of citations: A scientometric analysis of Lewontin, 1972

**DOI:** 10.1101/2021.09.19.460985

**Authors:** Jedidiah Carlson, Kelley Harris

## Abstract

“The Apportionment of Human Diversity” (1972) is the most highly cited research article published by geneticist Richard Lewontin in his career. This study’s primary result—that most genetic diversity in humans can be accounted for by within-population differences, not between-population differences—along with Lewontin’s outspoken, politically-charged interpretations thereof, has become foundational to the scientific and cultural discourse pertaining to human genetic variation. The article has an unusual bibliometric trajectory in that it is much more salient in the bibliographic record today compared to the first 20 years after its publication. Here, we show how the paper’s fame was shaped by four factors: 1) citations in influential publications across several disciplines; 2) Lewontin’s own popular books and media appearances; 3) the renaissance of population genetics research of the early 1990s; and 4) the serendipitous collision of scientific progress, influential books/papers, and heated controversies in the year 1994. We conclude with an analysis of Twitter data to characterize the communities and conversations that continue to keep this study at the epicenter of discussions about race and genetics, prompting new challenges for scientists who have inherited Lewontin’s legacy.

## Introduction

Richard Lewontin’s 1972 study, “The Apportionment of Human Diversity” [1] (hereafter referred to as “Lewontin 1972”), published as a contributed chapter in volume 6 of the book series, “Evolutionary Biology” [2], is widely considered to be a landmark publication in human population genetics research [3–5]. This paper is widely attributed as the originator of an aphoristic sound bite that is foundational to our understanding of human genetic diversity—”there is more genetic variation within populations than between populations”—variations of which have been colloquially cited far beyond the measurable scope of citations in the academic literature, ranging from educational materials developed by the National Human Genome Research Institute [6] to public television documentaries [7] to numerous references on contemporary social media platforms. Furthermore, Lewontin’s blunt interpretation of his results (“*Human racial classification is of no social value and is positively destructive of social and human relations. Since such racial classification is now seen to be of virtually no genetic or taxonomic significance either, no justification can be offered for its continuance*” [1]) has come to be ubiquitously associated (and in some cases, incorrectly credited) with the widely-held consensus that race is a social construct, not a biological one [8]. The far-reaching influence of Lewontin 1972 is partially evident in its bibliometrics: over the last half-century, it has amassed thousands of literature citations across a diverse range of disciplines; not only within Lewontin’s home fields of genetics and evolutionary biology, but also anthropology, medicine [9], psychology [10], sociology [11], and information science [12], many of which have gone on to become highly influential studies within their respective fields.

Though care must be taken to avoid conflating scientometric indicators with more subjective definitions of a publication’s impact and influence [13–15], a careful examination of the scientometric data surrounding Lewontin’s paper is useful in helping us formulate several pertinent questions about the paper’s history: What factors drove the extensive academic and cultural attention surrounding the paper, and how has that attention evolved over the last 50 years? How did the paper come to be associated with the “more genetic variation within populations than between populations” sound bite? Even though Lewontin’s contemporaries published several topically similar papers in the same era, why did Lewontin 1972 emerge as the most iconic? How has social media perpetuated and mutated the discourse surrounding this paper? Answering these questions may, in turn, help address a number of queries relevant to today’s researchers: what makes an ordinary research paper have an extraordinary bibliographic and/or cultural impact? How can scientists engage with socially sensitive research topics while maintaining their personal moral and ethical convictions? What are the benefits and landmines of using social media to communicate the results and implications of such research?

In the spirit of Lewontin himself, who constantly urged his colleagues to acknowledge the interpenetration between scientific research and the socio-ecological systems in which that research is embedded [16], we attempt to address these questions by examining patterns in the citing literature, major events surrounding the broader field of human genetics research, and Lewontin’s own career trajectory. We identify four factors that appear to have propelled Lewontin 1972 to its current iconic status: 1) citations in several highly influential books and papers, beginning almost immediately after its publication; 2) Lewontin’s influence through his popular books and media appearances in which he reiterated the results of his paper; 3) rapid technical advancements in molecular genetics in the early 1990s that would prompt and enable a new generation of population geneticists to revisit landmark studies from the 1960s and 1970s; and 4) several influential and/or provocative publications and events that coincided, mostly by chance, in the year 1994.

To better understand the ongoing cultural impacts of Lewontin 1972, we proceed to explore how the publication was referenced on Twitter during a 9-month period from 2020-2021 (coincidentally ending our data collection just a month before Lewontin passed away in July 2021). We found that direct references to the paper on social media are nearly non-existent, undermining the utility of standard altmetric indicators like the Altmetric Attention Score. However, an expansive corpus of tweets indirectly referencing Lewontin 1972 (which we term “dark citations,” following [17]) reveals that the concepts Lewontin presented 50 years ago continue to maintain a foothold in the cultural zeitgeist. We conclude with a discussion of how these colloquial online citations differ from the traditional bibliometric record, and how social media has, for better or worse, democratized conversations about population genetics research.

### The Bibliometric Trajectory of Lewontin 1972

As of June 15, 2021, Lewontin 1972 had received 3,076 citations according to Google Scholar and 1,991 citations according to Semantic Scholar, translating to an average rate of 39.8-61.5 citations per year. According to Google Scholar, Lewontin’s only published works with more citations are the opinion piece “The spandrels of San Marco and the Panglossian paradigm: a critique of the adaptationist programme,” coauthored with Stephen Jay Gould in 1979 [18], and two popular science books: *The Genetic Basis of Evolutionary Change* [19] and *Not in our Genes: Biology, Ideology, and Human Nature* [20]. The citation trajectory of Lewontin 1972 over time appears to be highly unusual, with only 15% of citations occurring in the first 30 years of the paper’s lifespan and the remaining 85% of citations occurring since 2002 (**Fig. 1a**). The distribution of the number of citations per year is roughly bimodal, with an initial weak pulse of citations peaking in the early 1980s and tapering off to nearly 0 citations per year by the end of that decade, a pattern that was first observed by Ruvolo and Seielstad in 2001 [3]. A second, much stronger, pulse of citations emerged in the early 1990s, jumped dramatically in the early 2000s, and grew steadily until the annual citation rate peaked around 2010-2015 (**Fig. 1b**).

**Fig. 1.**
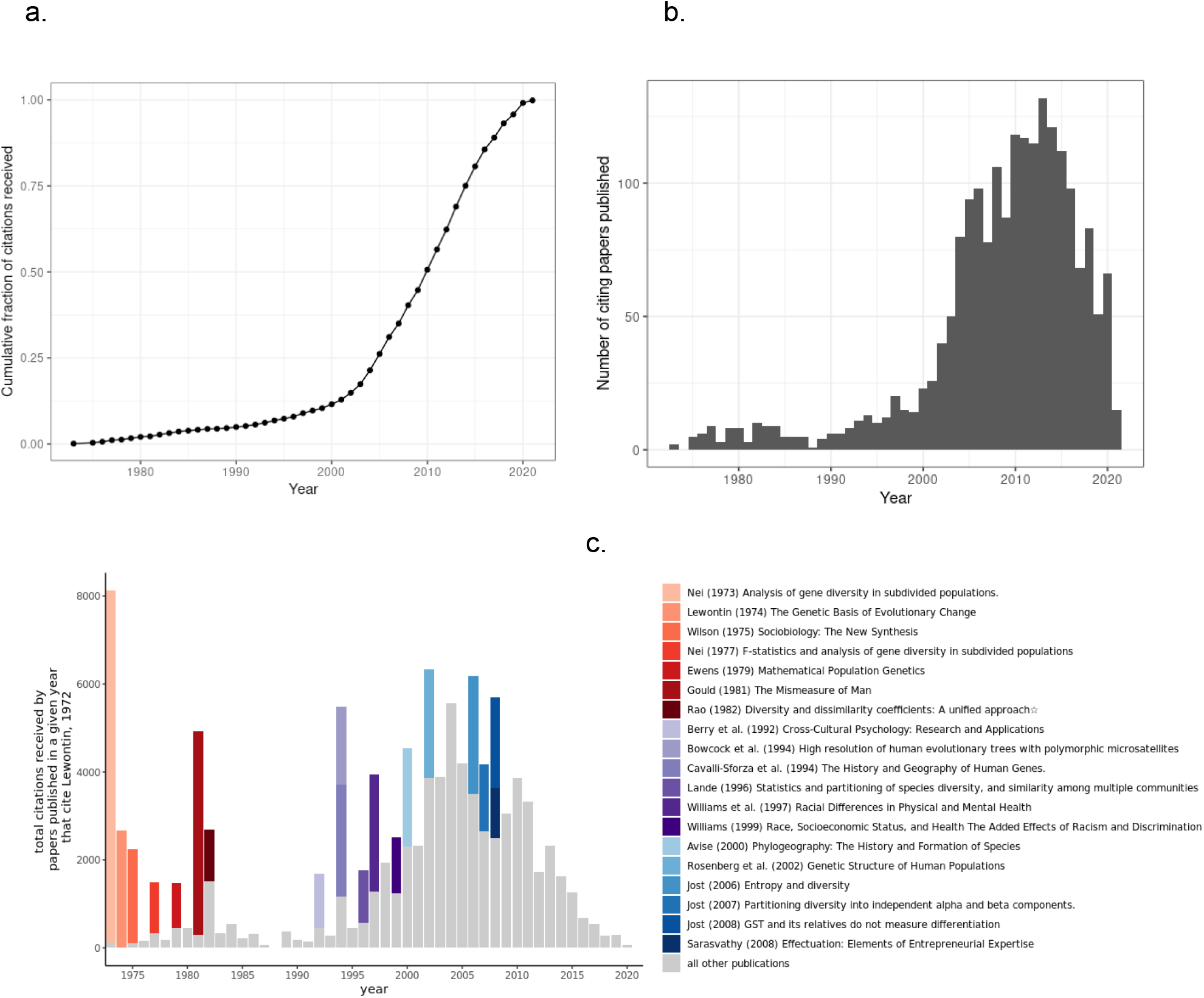
Bibliometric summary of Lewontin, 1972. a) cumulative distribution of citations over time; b) histogram of citations per year; c) histogram of 2nd-degree citations (i.e., among citing articles in a given year, the total number of citations they have received to date) over time. Contributions from the most highly influential papers (articles that went on to receive >1,000 citations, according to Semantic Scholar) that cited Lewontin 1972 are indicated in colored bar segments.

Ruvolo and Seielstad proposed two hypotheses for why Lewontin’s article received so few citations early on: either 1) the scientific community had “already come to believe that human races were effectively a scientific nonissue,” rendering the results of Lewontin 1972 “obvious,” or 2) the scientific study of race and genetics was “too politically charged” for further investigation [3]. Ruvolo and Seilstad effectively reject the former of their hypotheses in the very first paragraph of their paper, where they state:

*[Lewontin’s] findings surprised those who read the paper. Although typological notions of race had been on the decline in anthropology, many scientists and laypeople continued (and a few still continue) to expect substantial genetic differences between the groups they seemed able to recognize visually* [3].

Their second hypothesis, that the study of race and genetics was considered too politically taboo in the 1970s-1980s, is also easy to reject, in light of the numerous population genetics studies during that era that probed the topic of human genetic diversity within and between races [21–31]. These papers reflected an intense debate within the population genetics community: on one side, Lewontin, Latter, Nei and Roychoudhury had demonstrated that more genetic variation could be attributed to within-population differences than between-population differences (with Lewontin prominently proclaiming this as evidence that phenotypically-defined human races corresponding to those population groupings are not taxonomically significant), whereas Smouse, Spielman, and Mitton had demonstrated that the genetic markers that *do* differ between populations, although reflecting a small fraction of the overall genetic variation, could still be used to classify individuals into discrete groupings with some degree of accuracy (thereby arguing that phenotypically-defined human races corresponding to those population groupings *were* taxonomically significant to a degree) [5]. On the fringes of this debate were works by Nei and Roychoudhury, who repeatedly confirmed Lewontin’s finding of more variation within populations, but also maintained that the minimal between-population variance justified “the existence of a biological basis for the classification of human races” [31]. The bibliographic manifestation of this debate appeared to terminate rather abruptly in the early 1980s with a stalemate: two papers, Nei and Roychoudhury’s, “Genetic relationship and evolution of human races” (1982) [31] and Ryman et al.’s “Differences in the Relative Distribution of Human Gene Diversity between Electrophoretic and Red and White Cell Antigen Loci” (1983) [32] confirmed the earlier variance partitioning results, whereas Smouse et al. maintained that a multi-locus approach could be used to classify into discrete racial categories [30]. This historical context raises two key questions about the citation trajectory of Lewontin 1972: First, what are the factors that caused an uptick in citations in the early 1990s after a prolonged period of bibliographic stagnation? Second, why did Lewontin 1972 ultimately emerge as the most iconic study of human genetic diversity from this era?

## The Influence of Influential Citations

We first considered the extent to which the citation trajectory of Lewontin 1972 was shaped by having accumulated citations in other widely-cited scientific publications, which may have exponentially increased its exposure to other researchers. We identified 11 highly influential papers and 8 highly influential books/textbooks (we operationally define a publication as “influential” if it has received over 1,000 citations to date according to Semantic Scholar) that have cited Lewontin 1972. Seven of these were published in the 1972-1982 period, when there was an initial flurry of activity surrounding the topics of variance-partitioning and classification in human population genetics; notably all of these focused on reiterating either Lewontin’s results or his theoretical and statistical insights rather than his interpretations thereof [19,33–38] (**Fig. 1c**). Most notably, the influential citing publications from this era include three widely-read books that also set the stage for Lewontin’s prolonged feud with E.O. Wilson over sociobiology [39]: Lewontin’s own *The Genetic Basis of Evolutionary Change*, an edited compilation of lecture notes, published in 1974 [19], Wilson’s *Sociobiology: A New Synthesis* (1975) [34], and Gould’s *The Mismeasure of Man* (1982) [38] (note that the citation counts of these works sometimes drastically differ between Semantic Scholar and Google Scholar. For example, *The Mismeasure of Man* has been cited over 14,000 times according to Google Scholar but only ∼4,000 times according to Semantic Scholar).

No bibliographically influential citing publications appeared again until 1992, with Berry et al.’s psychology textbook *Cross-cultural psychology: Research and applications* [10]. Then in 1994, Bowcock et al. replicated Lewontin’s finding that most genetic variation exists within populations, using newly-available polymorphic microsatellite data [40]. The same year, Cavalli-Sforza et al. published their landmark book, *The History and Geography of Human Genes* [41]. In 1996, Russell Lande’s paper, “Statistics and partitioning of species diversity, and similarity among multiple communities,” again cited Lewontin 1972 for its contributions towards establishing a theoretical framework of variance partitioning within a species [42]. The next two influential citing articles were published in 1997 and 1999 by the eminent sociologist and scholar of health disparities, David Williams [11,43]. Seven other highly influential citing publications emerged during the 2000s, again turning the spotlight onto the technical aspects of Lewontin 1972 [12,44–48].

Most of these influential citations were unsurprising, in that they generally focused on building upon and reassessing Lewontin’s methods or reiterating his empirical results. The influential citations from Williams in 1997 and 1999, however, were an anomaly, not only because they came from the social sciences, but also because they shifted the focus onto Lewontin’s interpretations, paraphrased as follows: “*race is a gross indicator of distinctive social and individual histories and not a measure of biological distinctiveness*” [43]. While surveying the citing literature from the social sciences and the humanities, we came across another noteworthy paper, legal scholar Ian Haney López’s “The social construction of race: Some observations on illusion, fabrication, and choice,” published in the Harvard Civil Rights-Civil Liberties Law Review in 1994 [49]. This paper was reprinted a year later in the book *Critical Race Theory* [50], which is regarded as one of the foundational texts on the subject and has been cited over 3,000 times, yet neither Google Scholar nor Semantic Scholar had indexed this book as having cited Lewontin 1972. In light of these citations, we speculated that Lewontin’s provocative conclusions may have been a unique feature of his paper that particularly appealed to other scholars in the social sciences and humanities, contributing to its iconic status.

To further quantify the importance of citations from the social sciences, we annotated each citing journal article with the inferred academic discipline based on the Scopus journal taxonomy (limited to *N* = 1,330 citing articles for which these data could be retrieved) and examined the changes in relative abundance of each discipline among the citing articles over time. Although the majority of citing articles were published in life sciences journals over the paper’s lifespan (**Fig. 2a-b**), we found that citing articles in social sciences journals did have the highest relative prevalence between 1985-1990 (**Fig. 2b**), lending some credence to our hypothesis that the social sciences provided a boost in bibliographic exposure to Lewontin 1972 at a time when citations in the life sciences literature were particularly sparse. Citing articles in the social sciences, however, never grew to be cumulatively more common than those in the life sciences, and it was clearly in the life sciences where Lewontin 1972 saw the biggest surge in citations in the last 20 years (**Fig. 2a**). Even so, it is worth noting that references in the social sciences tended to differ in *how* they cited Lewontin 1972. Using Semantic Scholar’s “citation intent” flag, which infers whether a given citation was part of the background, methodology, or results, we found that citing articles in the social sciences almost always (82% of papers where data are available) cited Lewontin 1972 as background, compared to citing articles in the life sciences, in which only 37.5% cited Lewontin 1972 as background and the majority (62.5%) of citations were found in the methods and/or results.

**Fig 2.**
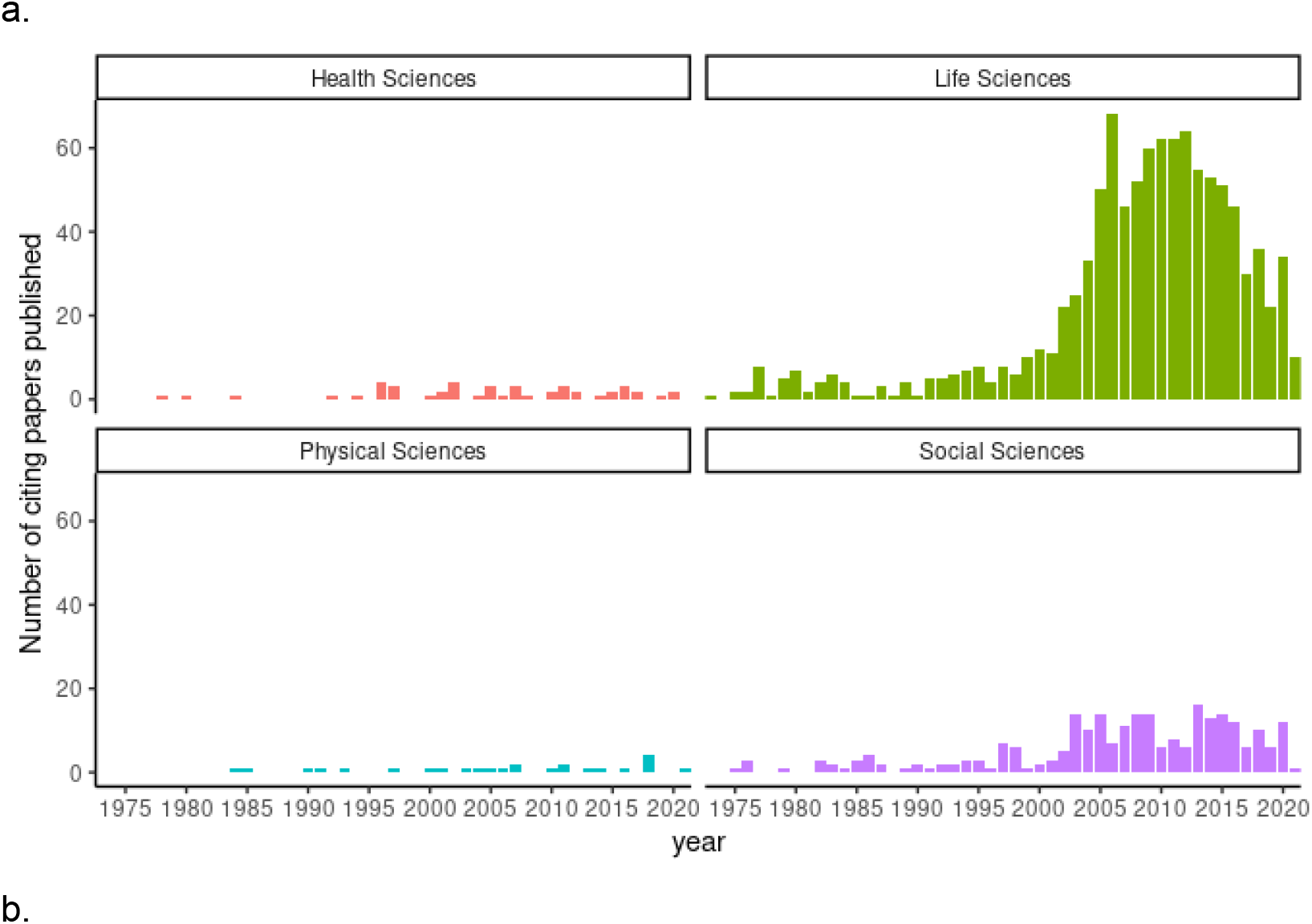
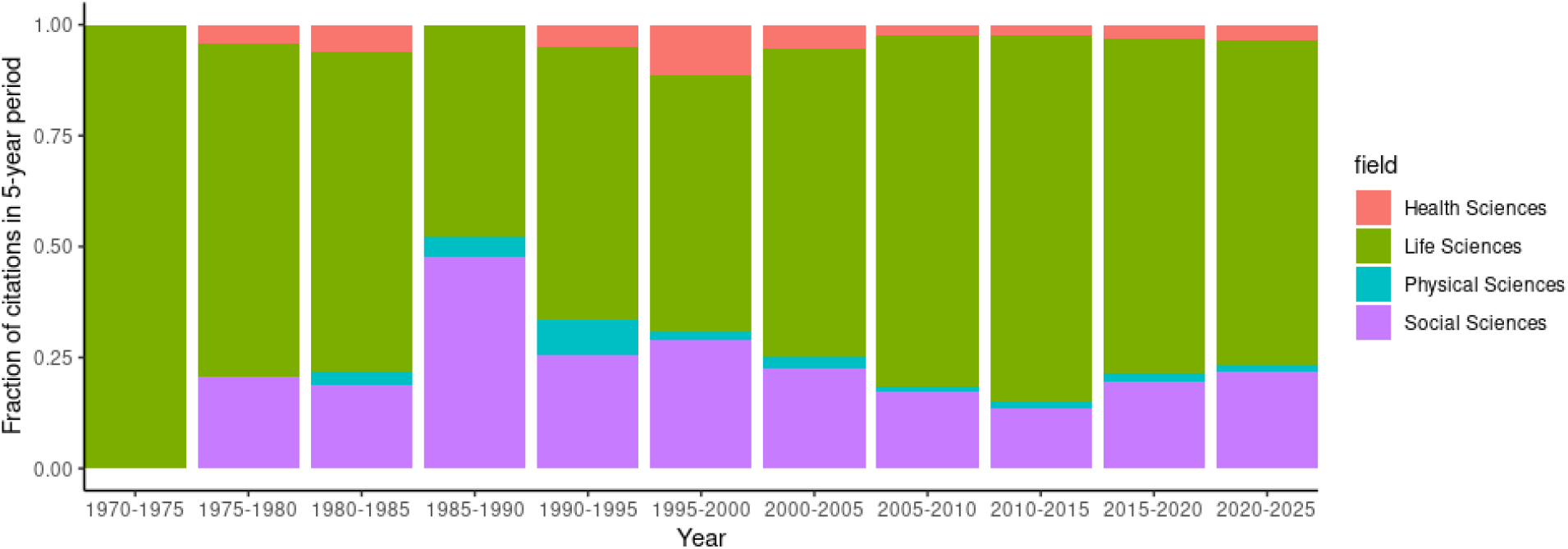
Breakdown of citation patterns for Lewontin, 1972 according to the inferred research domain of the citing articles. a) total number of citing articles in a given year, stratified by research domain; b) fraction of citing articles published in a given 5-year period within each of the four research domains.

### The influence of Lewontin’s popularity as a science communicator

Given that Lewontin’s *The Genetic Basis of Evolutionary Change* [19] was among the earliest influential citations, we also explored whether Lewontin had a tendency to self-cite his 1972 paper in his later works. We found only three other instances of self-citation, all published much later in his career: Lewontin and Hartl (1991), where the authors discuss the usage of population genetics in forensic DNA typing [51] (333 citations); Feldman and Lewontin (2008), in a chapter titled “Race, Ancestry, and Medicine” contributed to the book “Revisiting Race in a Genomic Age” [52] (54 citations); and Fujimura et al. (2014), that argued the authors of a 2012 paper in the journal *Sociological Theory* had made claims “based on fundamentally flawed interpretations of current genetic research” [53] (67 citations). Although the relative paucity of self-citations for this paper indicate that Lewontin mostly avoided the subject in his subsequent research (later in his career, he supposedly stated that he viewed the topic of genetics and race as “completely uninteresting” [3]), the results and conclusions of Lewontin 1972 were undoubtedly foundational to his interpretation of human genetic diversity and its relevance to social and political issues. For example, in *The Genetic Basis of Evolutionary Change* (1974), Lewontin forcefully reiterates his position:

*The taxonomic division of the human species into races places a completely disproportionate emphasis on a very small fraction of the total of human diversity. That scientists as well as nonscientists nevertheless continue to emphasize these genetically minor differences and find new ‘scientific’ justifications for doing so is an indication of the power of socioeconomically based ideology over the supposed objectivity of knowledge*. [19]

It is also worth noting that during the 1982-1994 period when the bibliographic influence of Lewontin 1972 appeared to have stagnated, Lewontin published four of his most celebrated books, all of which touch on the topic of race and genetics and Lewontin’s broader arguments against genetic essentialism, yet do not directly reference Lewontin 1972: *Human Diversity* (1982) [54], *Not in Our Genes* (1984) [20], *The Dialectical Biologist* (1985) [16], and *Biology as Ideology: The Doctrine of DNA* (1992) [55]. Lewontin also defended the conclusions of his paper in other public-facing media outlets throughout his career, including televised interviews in 1975 [56] (an episode of PBS’ NOVA program in which the famous physicist Richard Feynman was also profiled) and in 2003 [7]. Lewontin’s books, radio/television appearances, and his central role in the sensationalized debate over sociobiology in the 1970s-1980s may have served to keep Lewontin’s earlier scholarly work salient in the minds of his peers and the next generation of academics.

### Co-citation analysis

To search for additional clues about what may have sparked the influx of citations in the 1990s, we next examined which other papers were most commonly cited in the corpus of research articles that referenced Lewontin 1972 (**Fig. 3a**). The most frequent co-citation by far—including every year since 2009—was Nei’s “Analysis of gene diversity in subdivided populations” (1973) [33], which was also the most influential citing article (**Fig. 1c**). In addition, 3 of the 15 most common co-citations are also celebrated population genetics papers from the 1960s and 1970s: Kimura and Crow’s “The number of alleles that can be maintained in a finite population” (1964) [57], Nei’s “Genetic distance between populations” (1972) [58], and Nei’s “Estimation of average heterozygosity and genetic distance from a small number of individuals” (1978) [59]. When we compared the citation trajectories of these four papers, we found that they closely mirrored that of Lewontin 1972 (**Fig. 3b**), which implies that the underlying factors that contributed most strongly to the secondary surge in citations of Lewontin 1972 were likely common to other important population genetics papers from that era.

**Fig. 3.**
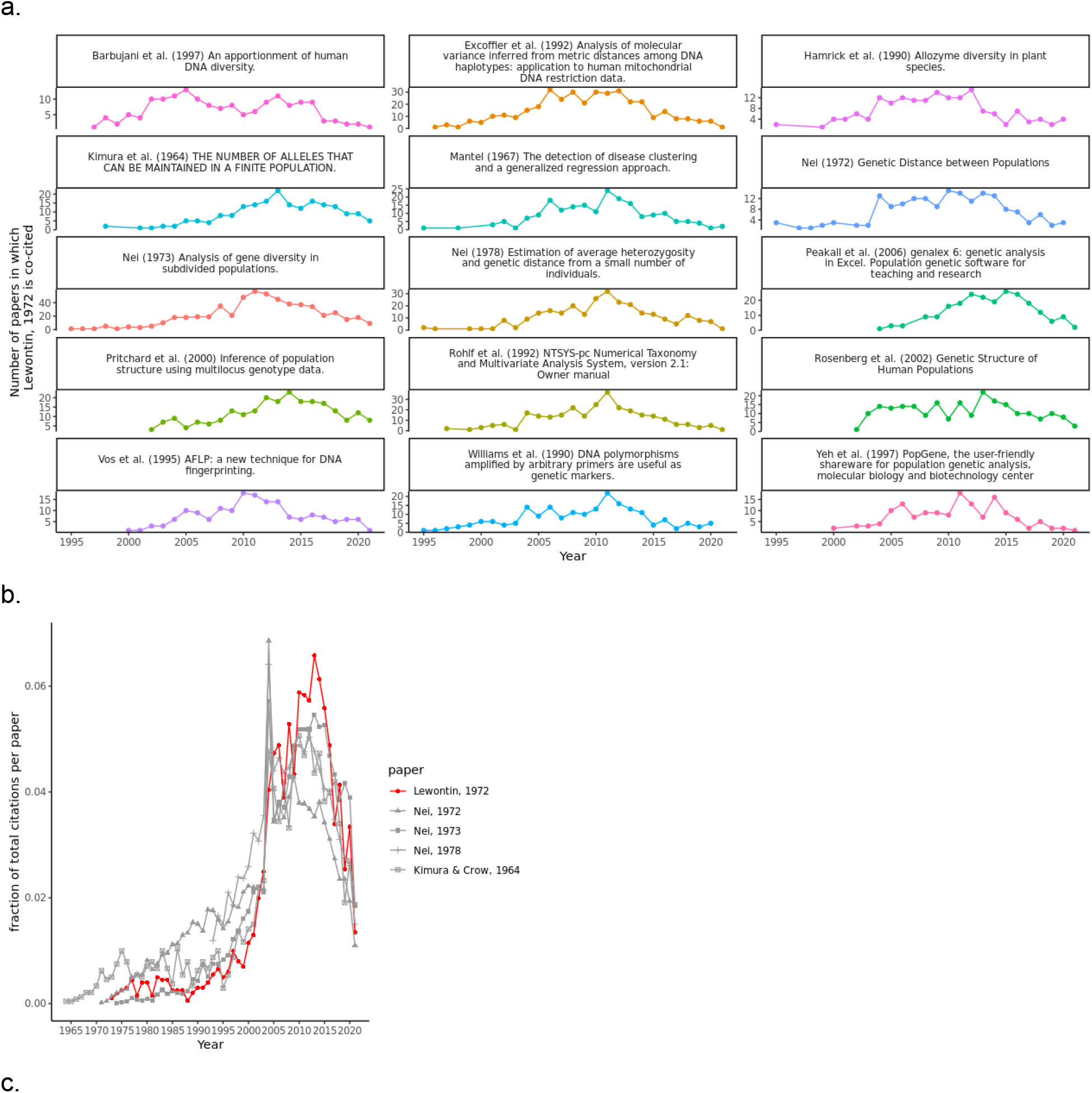
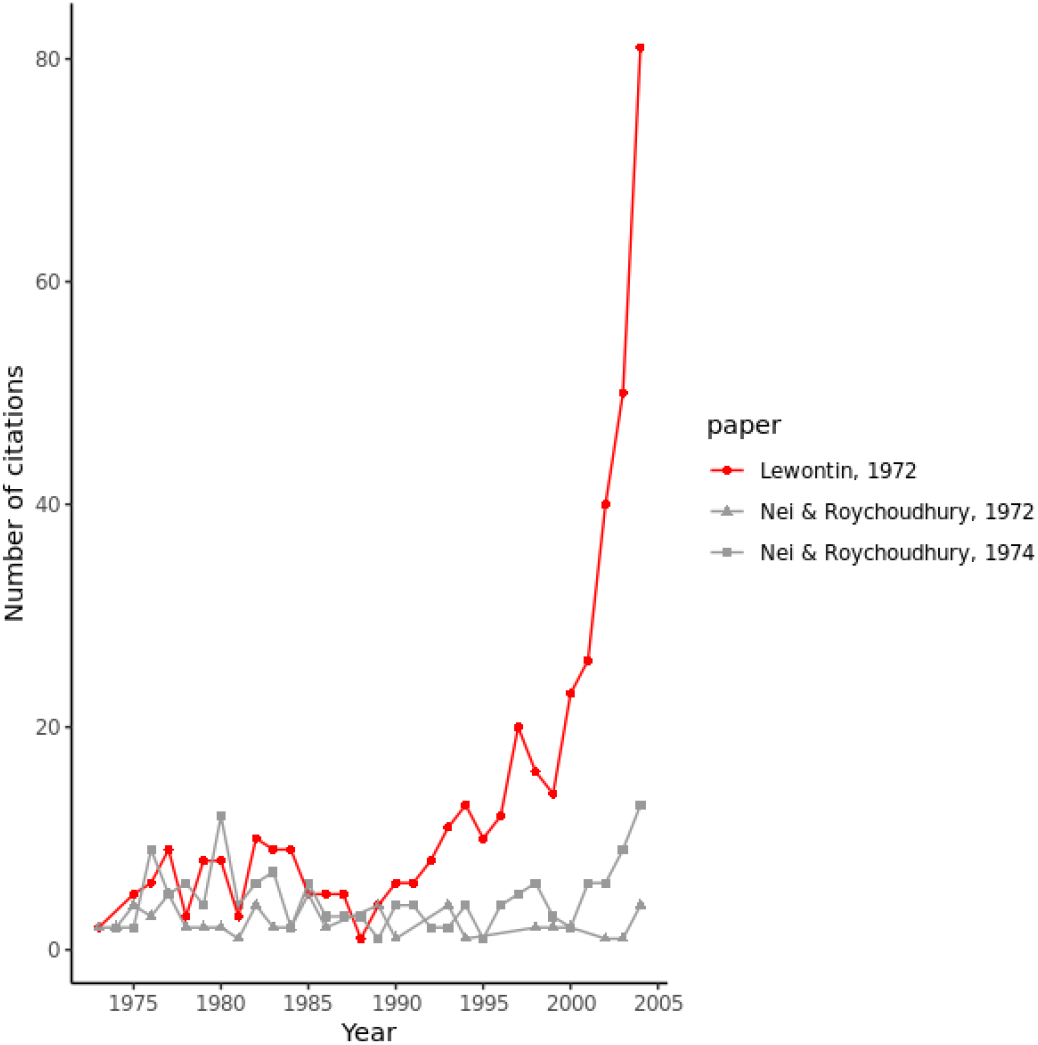
a) Co-citation frequencies over time for the 15 papers most commonly referenced alongside Lewontin, 1972. Data prior to 1995 are not shown due to much lower overall citations of Lewontin 1972. b) Citation trajectories of landmark population genetics papers (each published before 1980) that were among the most common co-citations of Lewontin, 1972. c) The citation trajectory of Lewontin 1972 compared to topically similar papers published by Nei and Roychoudhury in 1972 and 1974 that reported similar results.

As described in the previous section, one of the first influential citing papers in the 1990s was Bowcock et al., who, in 1994, replicated Lewontin’s variance-partitioning results using a novel type of genetic marker, polymorphic microsatellites [40] (another publication in 1994 would do the same using restriction fragment length polymorphisms [RFLPs] [60], though this paper has been less widely cited). Our co-citation analysis revealed that Lewontin 1972 was frequently cited alongside other articles describing novel genetic markers that emerged in the 1990s: Williams et al.’s “DNA polymorphisms amplified by arbitrary primers are useful as genetic markers” (1990) [61] which introduced a genotyping technique based on Random Amplified Polymorphic DNA (RAPD) markers, and Vos et al.’s “AFLP [Amplified Fragment Length Polymorphisms]: a new technique for DNA fingerprinting” (1995) [62] (**Fig. 3a**). When first introduced in 1990, RAPD markers offered several advantages over the prevailing source of polymorphic data, RFLPs, and were later described as being particularly useful in advancing empirical population genetics research [63]. This suggests that the renewed interest in Lewontin 1972 (and other papers from that era) was driven by the rapid evolution of new genotyping technologies in the late 1980s and early 1990s. This is all but confirmed by several of the remaining articles we found to be most frequently co-cited with Lewontin 1972, which focused on the same question of quantifying genetic variation within and between populations: Excoffier et al. (1992) used mitochondrial RFLP data [64], Barbujani et al. (1997) used autosomal RFLP data [65], and Rosenberg et al. (2002) used autosomal microsatellite data [45] (**Fig. 3a**). Other notable co-citations include several popular population genetic software programs first introduced in the 1990s and 2000s (genalex [66], NTSYS-pc [67], STRUCTURE [68], PopGene [69]), which suggests that rapid advances in computational power may have further precipitated and enabled interest in revisiting the early works of Lewontin, Nei, Kimura, and others using new datasets and computational methods. The other two top co-citations we identified were Mantel’s “The Detection of Disease Clustering and a Generalized Regression Approach” (1967) [70], which introduced a statistical test that came to be widely adopted in population genetics for evaluating population structure [71], and Hamrick and Godt’s “Allozyme diversity in plant species” (1990) [72], underscoring that the relevance of Lewontin’s work extended far beyond the human species.

### What happened to Nei and Roychoudhury?

One outstanding question that these citation and co-citation patterns still do not explain is why Lewontin’s paper went on to amass thousands of citations in the last 30 years, yet the comparable studies published by Nei and Roychoudhury in 1972 [21], 1974 [73], and 1982 [31] did not. Though the papers of Nei and Roychoudhury never received enough citations to be counted among the most frequently co-cited with Lewontin 1972, we confirmed that they are co-cited in several of the most highly-cited publications we identified, including Nei (1973) [33], Bowcock et al. (1994) [40], Cavalli-Sforza et al. (1994) [41], and Haney López (1994) [49]. In addition, the citation trajectories of Nei’s and Roychoudhury’s papers from 1972 and 1974 are very similar to that of Lewontin until the early 1990s (**Fig. 3c**). Furthermore, as early as the mid-1980s (and perhaps even earlier), both Lewontin 1972 and Nei’s and Roychoudhury’s work had been invoked in arguments rejecting the biological basis of race [39,74]. These bibliographic breadcrumbs suggest that for the first 20 years, Lewontin 1972 did not have broader exposure to fields outside of population genetics, nor was it viewed as having bibliographic precedence in the literature, nor did it occupy an exclusive role in shaping the scientific consensus that race is not taxonomically meaningful. However, the citation trajectories shown in **Fig. 3c** indicate there was an inflection point in the early 1990s when citing Nei and Roychoudhury fell out of favor, and indeed, only two of the 9 influential papers published after 1994 cited both Lewontin 1972 and Nei and Roychoudhury.

To resolve this question, we turned to several key papers in the corpus of citing literature that contrasted the rhetoric of Lewontin 1972 with that of Nei’s and Roychoudhury’s publications. As described by Haney López in 1994, the two diverged dramatically in their interpretations of the variance-partitioning results:

*Lewontin argued that biologists should abandon all talk of biological races […]. Nei and Roychoudhury agree that talk of biological races should be abandoned, but point out that there remain statistically significant differences between smaller population groups that justify the continued scientific division of humans by gene type* [49].

Brown and Armelagos would later characterize the conclusions of Nei and Roychoudhury to be logically inconsistent with their results:

*Interestingly, despite these very low figures [of between-population variance], [Nei and Roychoudhury] went on to discuss ‘the pattern of evolution of the three major races’ (p. 11). This speaks to the logical disconnect shown by many researchers who simultaneously prove the irrelevance of genetic race and then proceed to discuss the genetic evolution of races* [4].

In Stephanie Malia Fullerton’s retrospective analysis [75] of Kwame Anthony Appiah’s celebrated essay on the philosophy of race, “The Uncompleted Argument: Du Bois and the Illusion of Race” (1985) [74], she explains how Appiah attributes Nei and Roychoudhury (1982) as paramount evidence for rejecting the biological basis for the concept of human races (in a footnote, Fullerton acknowledges Lewontin 1972 as being one of a handful of scientific papers that was “publicly important much earlier,” but does not elaborate on what differentiated Lewontin’s conclusions from those of Nei and Roychoudhury). Even so, Appiah was aware of the fact that Nei and Roychoudhury still essentially held the view that race was taxonomically meaningful:

*[Appiah] also acknowledged that the geneticists he cited [Nei and Roychoudhury] were those who “believe in human races” but disputed their claim that their data “shows the existence of a biological basis for the classification of human races” [75]*.

Haney López, who cites Appiah extensively, takes this framing of Nei and Roychoudhury as racialists even further, concluding that “Nei and Roychoudhury reflexively fall into the comfortable habit of White supremacy in science” [49]. These viewpoints suggest that by 1994, the variance partitioning results of Lewontin, Nei, and Roychoudhury had been vindicated as factually correct and widely accepted, but only Lewontin’s blunt interpretations thereof were viewed as being consistent with a multidisciplinary synthesis rejecting the biological basis of race, whereas Nei and Roychoudhury’s racialist views rendered them outdated and irrelevant.

### The impact of the Human Genome Project

The Human Genome Project (HGP) was one of the most massive scientific endeavors of the 20th century, and it catalyzed an extraordinary increase in the amount of data available for research on human genetic variation. Given the foundational importance of Lewontin’s work towards the interpretation of genomic data, we considered the possibility that the HGP’s architects were familiar with Lewontin 1972 and helped propel it into the spotlight. Chronologically, this seems like a plausible explanation: the HGP was announced in 1990 and the completion of the first drafts of the human genome were published in 2001; these two years roughly align with marked increases in citations of Lewontin 1972 (**Fig. 1b**). However, although planning documents for the HGP mentioned the importance of the project’s ethical, legal, and social impact (including how genomic data might be misused to “advance eugenics or prejudicial stereotypes” [76]), we found no evidence in the bibliographic record that Lewontin 1972 was ever explicitly referenced by the organizations and key personnel involved in initiating the HGP, nor was it referenced in any of the papers or commentaries appearing in special issues of *Nature* and *Science* announcing the completed draft sequences of the human genome in 2001.

Although we found no evidence that the HGP had a direct bibliographic connection to Lewontin 1972, it likely played a role in popularizing the sound bite “there is more genetic variation within populations than between populations,” which is deeply intertwined with Lewontin’s legacy today [8]. Earlier literature suggests Lewontin himself was circulating versions of this sound bite in popular media as early as the mid 1970s [39], but it seems that, over time, the aphorism came to be disconnected from Lewontin as its originator. This disconnect is plainly apparent in A.W.F. Edwards’ 2003 critique of Lewontin 1972 [77]. Edwards begins his paper by mentioning that when the first draft of the human genome was published in *Nature* in 2001, the print version of the journal shipped with a compact disc of educational materials pertaining to human genomics research (the materials from this CD, titled “Understanding the Human Genome Project,” are currently hosted online by the National Human Genome Research Institute [6]). These materials include a module about human genetic variation that stated “there is more genetic variation within populations than between populations,” but without citation. Edwards took issue with how this statement had been used to “play down the genetical differences among human populations…usually without reference” [77], but despite his personal and scholarly familiarity with Lewontin, he was entirely unaware of Lewontin’s past association with this statement [78]. Edwards’ eventual discovery that the statement could be traced back to Lewontin 1972 led him to effectively attribute the paper as an uncited source for both the HGP and a backlog of highly influential news media that had indirectly popularized the paper’s conclusions. One such example we identified was a New York Times article from 2000 in which the author briefly summarizes the within- and between-population variance partitioning results (which at this point had been replicated by multiple research groups) and contains quotes from several prominent scientists (including those involved with the HGP) indicating their familiarity with these results, but Lewontin is never mentioned [79].

Though Lewontin 1972 was being cited more than ever by the turn of the 21st century (**Fig. 1b**), these anecdotes suggest that the results of the study had become so widely known and accepted that attribution was not always considered necessary. Paradoxically, the HGP’s singular focus on a single genome had, from the onset, incited researchers to wonder how much was omitted from that narrow view of human genetics [80]. By giving scientists more of a reason to think about human genetic variation, the HGP may have prompted them to rediscover and bibliographically reassert Lewontin 1972 as they sought to augment our understanding of human diversity with data from more individuals and populations. Ironically, Lewontin was highly critical of the ideological motivations and rationale of the HGP and published several essays in the New York Review of Books, which were ultimately compiled in his 2000 book, “It Ain’t Necessarily So: The Dream of the Human Genome and Other Illusions” [81].

### Standard altmetric indicators of Lewontin 1972

Citations of Lewontin 1972 in the academic literature have declined precipitously in the last 5 years (**Fig. 1b**), rendering traditional bibliometrics relatively ineffectual for understanding the ongoing impacts of the paper. However, the recent rise of social media as a vehicle for discussing scholarly research has enabled new paradigms for documenting and measuring how papers are shared and discussed [82]. The Altmetric Attention Score, which aims to quantify the attention received by scholarly works on social media, news media, blogs, Wikipedia, and other non-traditional sources of citations, is one of the most popular altmetric indicators used by the research community. As of June 15, 2021, Lewontin’s 1972 paper had an Altmetric attention score of 85, with only 17 tweets from 17 unique users directly referencing the paper [83]. In contrast, Edwards’ 2003 critique had received nearly 10-fold more directly-referencing tweets (145 tweets from 102 unique users, as of June 15, 2021) and had a higher Altmetric Attention Score of 109 [84] despite having only received approximately 1/10th as many literature citations as Lewontin 1972 (318 citations of [77], compared to 3076 citations of [1], per Google Scholar). Although its relatively modest Altmetric Attention Score and lack of “indexable” Twitter citations (i.e., tweets including a DOI or URL link to an online version of the paper) might create the illusion that Lewontin 1972 has little impact within the rapidly changing landscape of social media, a closer examination shows that it is widely and actively discussed on Twitter on a daily basis, underscoring a prolonged cultural influence that cannot be directly ascertained from traditional bibliometrics or even standard altmetric indicators.

### The Impact of Lewontin 1972 as revealed by “dark citations” on Twitter

Using the Twitter API, we collected all tweets/retweets containing the word “Lewontin” that were posted over a 9-month period from 27 August, 2020 to 25 May, 2021, resulting in a collection of 2659 tweets/retweets, an average of 9.8 tweets/retweets per day during this period. We hypothesized that these tweets would contain a number of “dark citations’’ to Lewontin 1972, defined by [17] to be references to scholarly works that do not necessarily include traceable links such as DOIs or URLs. After excluding 152 tweets/retweets referencing the R.C. Lewontin Early Award from the Society for the Study of Evolution, which was soliciting nominations during this period, we confirmed this dataset did not contain any additional off-topic tweets, e.g., from (or in response to) unrelated users with the surname “Lewontin.” A timeline of these tweets and their retweets during this period is shown in **Fig. 4a**. This timeline paints a picture of a steady, ongoing conversation about Lewontin’s work rather than a flurry of activity surrounding any specific controversies or events. Most tweets (1636/2507; 65%) were original unique tweets, not retweets; moreover, the 2507 tweets/retweets came from 1589 unique users, demonstrating that these tweets are not simply the output of a few individuals with a particularly strong interest in Lewontin. Of the 1636 original tweets, 1381 were quote-tweets or replies to other tweets, suggesting that the majority of Twitter references to Lewontin primarily arise in debates and conversational contexts. Though most (1057/1636) of the original tweets were in English, the following languages each accounted for at least 10 tweets in our dataset: German, Spanish, French, Italian, Portuguese, and Turkish. Furthermore, we observed tweets in Arabic, Danish, Japanese, Norwegian, Russian, and Swedish, indicating a linguistically and likely geographically diverse audience. The most frequently used words present in this dataset of tweets are summarized as a wordcloud in **Fig. 4b**, which shows many of the most common words in this dataset refer to Lewontin’s other highly-cited works with coauthors: 512 include “Gould,” 269 include “Levins” and 91 include “Kamin” or “Rose,” referring to coauthors Stephen Jay Gould [18], Richard Levins [16], Steven Rose, and Leon Kamin [20].

**Fig. 4.**
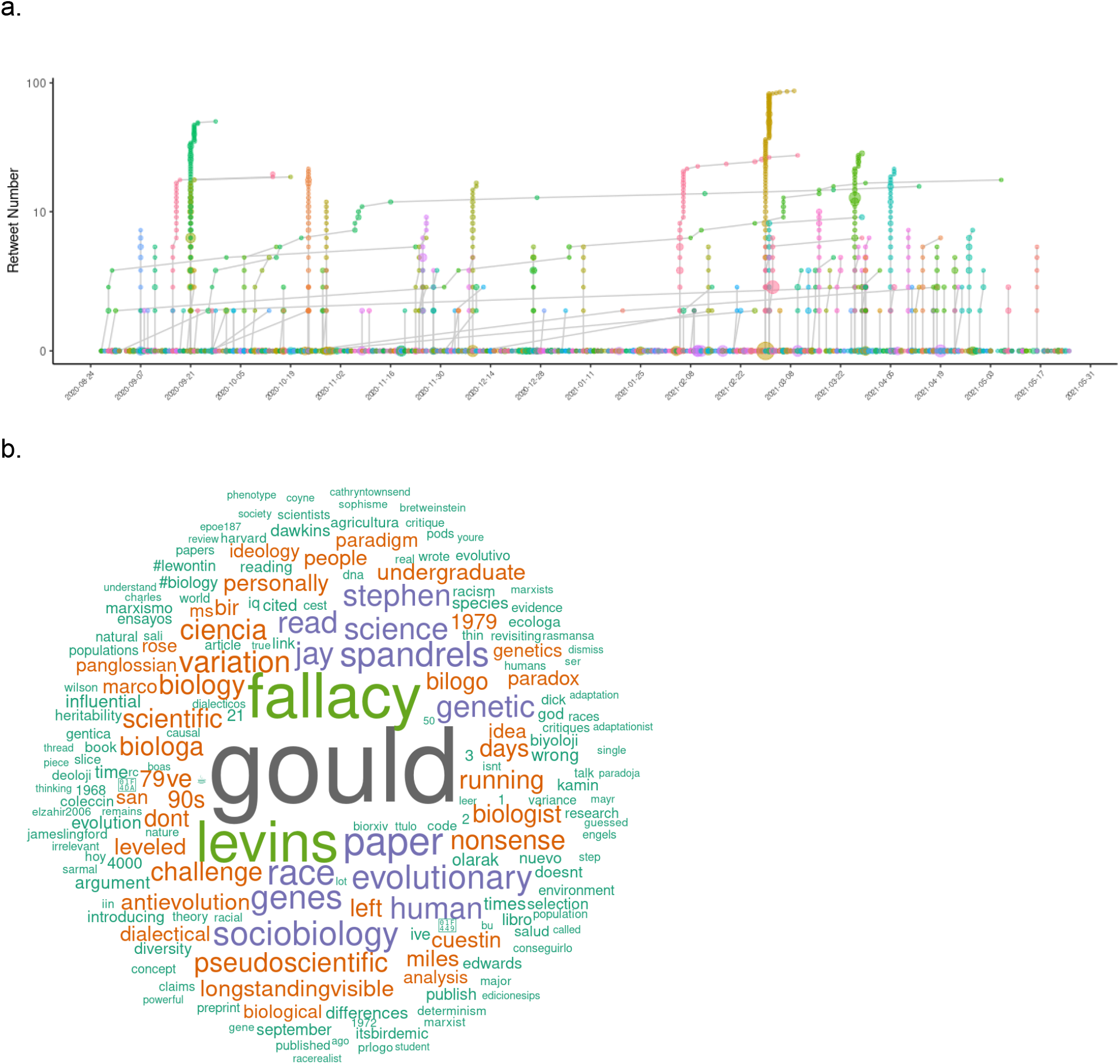
Summarizing 9 months of Twitter data for tweets referencing “Lewontin.” a) timeline of tweets and retweets in our dataset. Each original tweet is shown as a point along the x-axis. For tweets that were retweeted, the retweet trajectories are indicated by linked line segments growing along the y-axis as the number of retweets increases over time. The size of each point indicates the number of followers of the corresponding user. b) wordcloud representing the most frequent words found among the 2507 tweets collected (including retweets).

The most frequently used word in this dataset that is not a coauthor’s surname is “fallacy,” found in 10% (243) of the 2507 tweets/retweets. This refers to the phrase “Lewontin’s Fallacy,” popularized by Edwards’ critique of Lewontin 1972 [77]. A broader search for keywords relevant to Lewontin 1972 within these tweets revealed that 21.4% of the tweets (536/2507, or ∼2 tweets per day during the the 9-month data collection period) include the words “fallacy,” “diversity,” “race,” “racial,” “racism,” “racist,” “15%,” “85%,” or “variation” (we included “15%” and “85%” in these filtering criteria because they refer specifically to the percentages of human genetic diversity Lewontin attributed to between-race and between-individual differences, respectively [1]). In contrast, citations of Lewontin 1972 account for only 4% (3076/75637) of Lewontin’s total citations, according to Google Scholar. This suggests that Lewontin 1972 has an extraordinarily outsized influence in the social media ecosystem, relative to Lewontin’s broader body of work.

We also applied our recently-developed social media audience segmentation method [85] to categorize the users that engaged in conversations about Lewontin 1972 on Twitter over the 9 month data collection period. Briefly, we identified the followers of each of the 303 unique users whose tweets/retweets contained keywords indicating a specific reference to Lewontin 1972 (described above) then applied a statistical model to identify the most common co-occurring words in the bios of each focal user’s followers as an indicator of the network(s) each user is affiliated with. As described in [85], we interpret the properties inferred from each focal user’s followers as characteristic of that focal user, according to the principle of network homophily. The results of this audience segmentation are presented in **Fig. 5**. Based on the presence of particular keywords in the inferred audience topics (using the criteria described in [85]), we estimate that 44.2% of users in this dataset are primarily affiliated with academic communities, including medicine (sector 4), philosophy/psychology (sector 7), ecology (sector 8), genomics/bioinformatics (sectors 10 and 11) and evolutionary biology (sector 12), roughly mirroring the distribution of literature citations across life sciences, health sciences, and social sciences as portrayed in **Fig. 2** (though none of the academic audience sectors appear to align with fields in the physical sciences).

**Fig 5.**
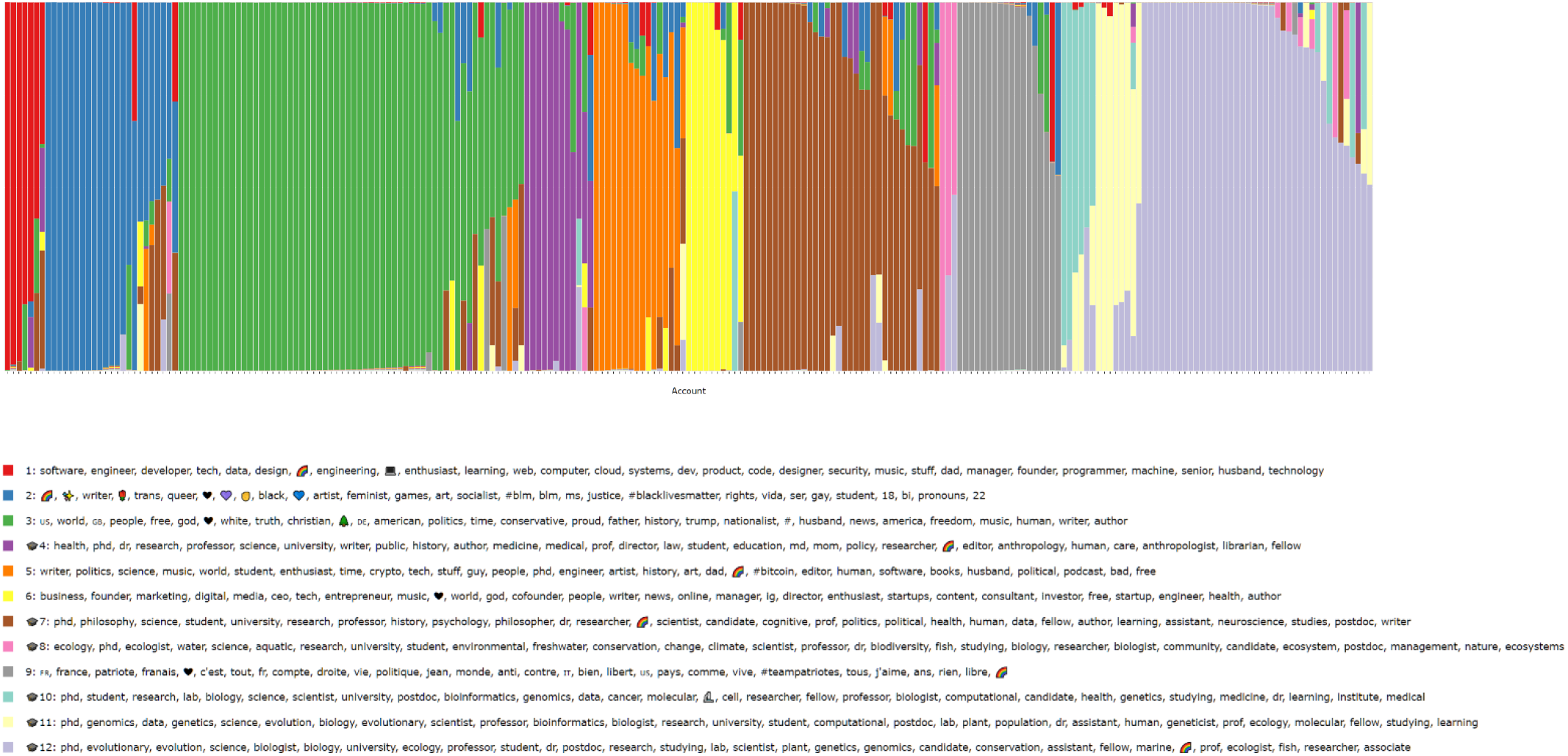
Twitter audience segmentation analysis for users indirectly referencing Lewontin 1972. Each unique user that tweeted about that paper is represented by a vertical stack of 12 colored bars that represent the user’s estimated membership probability in each of the 12 inferred audience sectors. The top 30 keywords, hashtags, or emoji associated with each audience sector are shown in the legend. Audience sectors inferred to correspond to academic communities are indicated with a 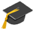 emoji at the beginning of the list of associated keywords.

Two audience sectors appeared to be primarily characterized by the industry/occupation of their constituent users: software engineering/development (sector 1) and business & marketing (sector 6), which together accounted for 7.4% of users in the dataset. The remaining audience sectors, accounting for 48.4% of users in the dataset, were primarily characterized by political ideology/affiliation. Most (52%) of these politically-affiliated users aligned with right-wing American politics (sector 3) and likely fell along a spectrum ranging from mainstream Republican supporters (evidenced by the keywords 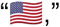 “conservative,” “trump”) all the way to adherents of far-right identitarian ideologies (evidenced by the keywords “white,” and “nationalist”). Sector 9 was mainly represented by French keywords and political terminology; the presence of the hashtag “#teampatriotes” in the top keywords (which appears to be popular among French twitter users who support far-right political candidates) and at least one tweet originating from a prominent French white nationalist (see Discussion) suggests that users associated with this sector also tend to align with far-right ideologies. Conversely, sector 2 appeared to capture users aligned along the spectrum of left-wing politics, evidenced by the keywords “socialist,” “feminist,” and “#blacklivesmatter,” accounting for approximately 20% of the politically-affiliated audience. The remaining audience sector, sector 5, appeared to be a “generalist” category of users interested in politics among a variety of other topics (including science, culture, technology, and history), but the top keywords do not indicate a specific political ideology and closer qualitative examination of the users affiliated with this sector suggests they identify across the political spectrum.

## Discussion

Our analysis of the bibliometric and altmetric indicators of Lewontin 1972 provides an important backdrop for understanding and contextualizing the significance of its scientific and cultural impacts. The citation trajectory shows that references to Lewontin 1972 were sustained at a modest rate through the 1970s and early 1980s (peaking at around 10 citations per year in 1982), then tapered off into the late 1980s, only to be followed by a surge of bibliographic attention beginning in the early 1990s and extending well into the postgenomic era, peaking at over 130 citations per year in 2013, over 40 years after its initial publication. The events and publications surrounding Lewontin 1972 that we analyzed here are summarized on a timeline in **Fig. 6**.

**Fig. 6.**
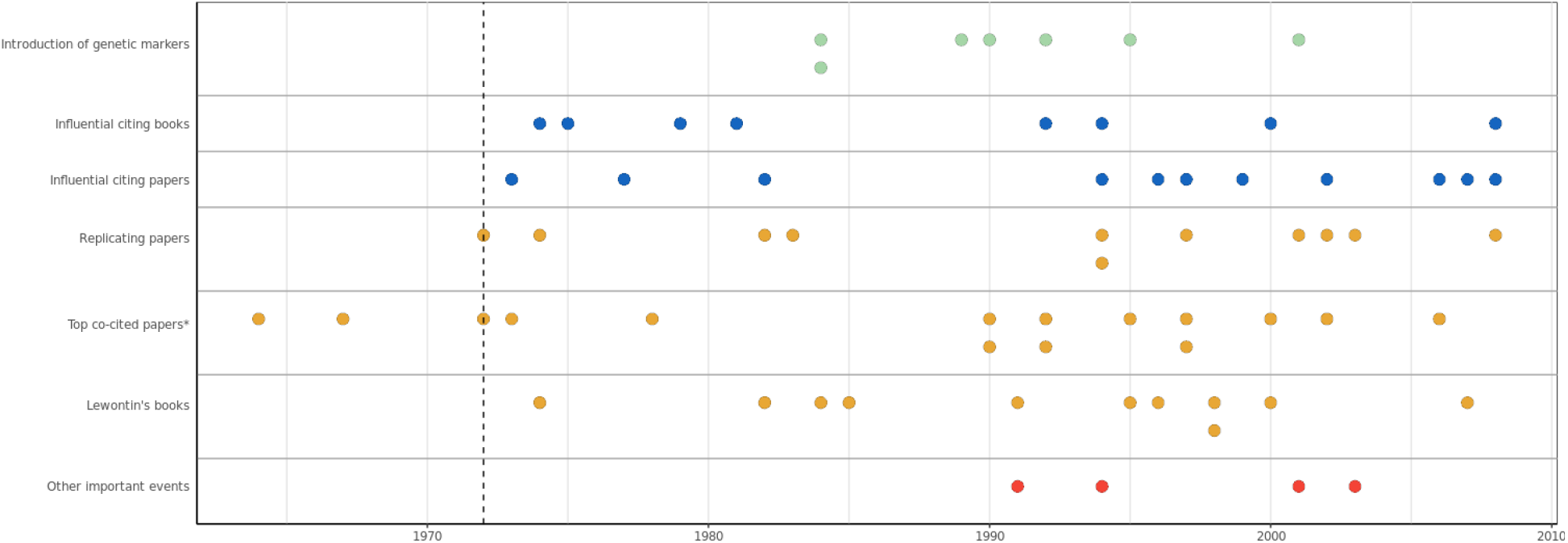
Timeline summarizing the relevant events, publications, citations, and co-citations surrounding and involving the bibliographic history of Lewontin 1972.

As revealed by our analysis of influential citing papers and co-citations, there were several trends and events that coalesced in the early 1990s (and in 1994, specifically), to propel Lewontin 1972 to its current iconic status. First, the field of empirical population genetics research entered into somewhat of a renaissance in the early 1990s [86]. By the early 1980s, the initial flurry of debate over Lewontin’s results had mostly subsided, and data from classical genetic markers (e.g., blood groups, proteins) had been extensively mined. A decade later, the emergence of novel genetic markers that could be easily assayed genome-wide (e.g., RFLPs, microsatellites), along with increasingly sophisticated computational tools for analyzing these data, made it possible to empirically revisit decades-old theories, hypotheses, and results in the field. With respect to Lewontin 1972, the fruits of this renaissance were first evident in two 1994 studies that replicated Lewontin’s results: Dean et al., which analyzed RFLP data [60], and Bowcock et al., which analyzed microsatellite data [40]. By this point, Lewontin 1972 was both well-established in the population genetics literature and increasingly recognized for its contributions to the scientific consensus rejecting the concept of biological races, in large part due to Lewontin’s books and media appearances throughout the 1970s and 1980s [39]. In the same year these replication studies were published, two other highly influential works would cite Lewontin 1972: Cavalli-Sforza et al. published their celebrated book, *The History and Geography of Human Genes* (later described as “the great synthesis of genetic data with historical, archaeological and linguistic information” [87]), and Haney López published a paper invoking Lewontin’s results as foundational to Critical Race Theory, which was republished in one of the most important books on the topic [50] the following year.

Entirely by coincidence, 1994 also marked the year that Richard Herrnstein and Charles Murray published their book, *The Bell Curve* [88], which immediately attracted widespread controversy over its claims that racial differences in IQ were due to innate genetic differences between races. Though Lewontin had sparred with Herrnstein decades earlier about the heritability of IQ [89] (and by this time had established himself as a popular outspoken opponent to the genetic essentialism espoused by Herrnstein and Murray [20]), Lewontin 1972 was not cited in *The Bell Curve*, and Lewontin mostly avoided engaging with this particular controversy, at least according to the bibliographic record. The results and conclusions of Lewontin 1972 were nonetheless repeatedly cited by critics of Herrnstein and Murray. For example, the specific citation of Lewontin 1972 in Gould’s *The Mismeasure of Man* is mentioned in a 1994 Washington Post letter to the editor refuting claims made in *The Bell Curve* [90] (two years later, *The Mismeasure of Man* was revised and republished with a new cover prominently proclaiming it was “the definitive refutation to the argument of *The Bell Curve*”). In 1995, philosopher Ned Block wrote a scathing critique of Herrnstein and Murray that mentions the results of Lewontin 1972 as being “widely accepted by all sides,” (note that Block cites Lewontin’s *Human Diversity* book as the source of these results rather than the 1972 paper) and describes Murray’s grasp of variance partitioning as “pathetically misunderstood” [91]. Lewontin’s relevance to this controversy is perhaps best summarized by Charles Murray himself, who, in a 2005 retrospective op-ed about the impact of his book, stated (incorrectly and somewhat obtusely) that “Richard Lewontin originated the idea of race as a social construct in 1972” [92]. Though we are unable to put any numbers to its effect, *The Bell Curve* undoubtedly played a role in popularizing Lewontin 1972 in the 1990s. Given that multiple scholars around this time were beginning to express their view that Nei and Roychoudhury’s conclusions were both logically inconsistent and tacitly supportive of racialist perspectives [4,39,49,74], we also speculate that *The Bell Curve* controversy may have fanned the flames of these arguments and significantly contributed to the ensuing bibliometric stagnation of Nei and Roychoudhury’s papers. The role of the Human Genome Project in bringing direct attention to Lewontin 1972 is tenuous at best—we found no evidence of Lewontin 1972 being explicitly cited in HGP-related publications and communications, despite the seemingly obvious role the HGP played in amplifying the “sound bite” of the paper.

Though literature citations of Lewontin 1972 have tapered off in the last five years, the Twitter data we analyzed showcases how the study has persisted in the academic and public discourse, accumulating roughly 2 tweets per day, with approximately equal attention from academic and non-academic audiences. Our analysis differs from most other altmetric studies because we primarily focus on “dark citations” found on social media (i.e., references to the study that do not directly link to a DOI or URL and thus are not tracked by Altmetric and other altmetrics data brokers). The concept of “dark citations” has been previously used to describe the practice of referencing prior published work in a journal article without an explicit citation [17], but it is perhaps even more relevant for altmetrics where much of the social media attention surrounding a paper takes place in colloquial threaded discussions. Such data are arguably a more salient indicator of a paper’s societal/cultural impact than standard bibliometric/altmetric indicators because they demonstrate the degree to which a paper’s results and conclusions have become embedded in the public mindset, to the extent that directly referencing the paper is superfluous. In the case of Lewontin 1972, our analysis shows how the paper continues to be embroiled in the cultural reckoning of defining and applying the concept of human races.

We acknowledge that our search criteria for these “dark citations” were not exhaustive; tweets containing some variation of the phrase “there is more genetic variation within populations than between populations” were excluded unless they explicitly included the word “Lewontin.” This may have biased our dataset to more thoroughly sample tweets that were critical of the study and/or Lewontin himself: because the phrase “Lewontin’s Fallacy” is popular among Lewontin’s critics and detractors, their tweets would have been included, whereas those who supportively repeat the “sound bite” of the paper may be entirely unaware of who Lewontin is and his role in popularizing this interpretation. Ironically, our data shows that, among non-academic audiences on social media, the groups most ideologically opposed to Lewontin’s claims play a significant role in maintaining the connection between Lewontin and the broader concept that “there is more genetic variation within a population than between populations.”

It should be noted that the tweets in our dataset that mention “Lewontin’s Fallacy” can be construed as indirect references to Edwards’ critique [77] rather than Lewontin’s original paper [1]. Though this distinction has no practical bearing on our analyses, it may be a useful one to make in understanding the motivations of those who continue to keep this paper in the spotlight. Many critical tweets in our dataset appear to invoke “Lewontin’s fallacy” simply as a rhetorical cudgel in an attempt to dismiss an opposing argument as logically invalid. For example, one Twitter user states: “This fallacy is so well known in science that it has a name: Lewontin’s fallacy.” Similar tautological claims are echoed by ideological critics in the scientific literature; for example, [93] states “[a fallacious claim that genetic similarity among humans negates phenotypic differences] is so common in the biological and social science literature that it even has its own name: *Lewontin’s fallacy*, named for a biologist who popularized it.” Such statements lionize Edwards’ critique as the authoritative interpretation of Lewontin’s results and paint Lewontin as a solitary proponent of this claim, despite more contemporary research that has largely vindicated Lewontin’s interpretation (e.g., [65,94–97]). Moreover, we find that the Twitter users who vehemently oppose the conclusions of Lewontin 1972 often have significant overlap with extreme far-right political communities, underscoring how rejection of Lewontin’s interpretation has become a tenet of white nationalist ideology.

One such individual who appeared in our dataset is Renaud Camus, a prominent white nationalist French writer who gained notoriety for coining the term “Great Replacement,” a conspiracy theory that postulates White European populations are being demographically and culturally replaced by non-White immigrants through policies enacted by “the global elites” [98]. Camus’ tweet states “*Jamais la Science ne se sera montrée plus serve*,” translating to “Science will never have shown itself to be more useful,” and quotes another user who wrote:

*“It was in fact in 1972 that the geneticist Richard Lewontin published the article ‘The Apportionment of Human Diversity’, which should be a landmark in the construction of anti-racist dogma*.*”* (Translated from French via Google Translate).

This conversation traces back to an earlier tweet posted by Camus, where he claims:

*“In 1976 the Haby law* [legislation introducing educational reforms in France], *inaugurat[ed] the single college, abolished classes, eradicated the cultivated class and therefore culture; at the same time the dogma of the inexistence of races eradicated the white race and therefore Western civilization*.*”* (Translated from French via Google Translate).

Taken in the context of this thread, we interpret Camus’ original tweet to insinuate that he views Lewontin as the architect of an “anti-racist dogma” that put into motion the “eradication” of the white race (not unlike Charles Murray’s incorrect attribution of Lewontin as the architect of “race as a social construct” [92]). Given that Camus’ Great Replacement theory is widely regarded as a core belief among white nationalists [99] and has been explicitly cited in manifestos written by the perpetrators of mass shootings in Christchurch, New Zealand, Pittsburg, Pennsylvania and El Paso, Texas [100], his familiarity with Lewontin’s paper is an alarming reminder that scientific research does not occur in a political vacuum. Just as Lewontin was acutely aware of the political minefield surrounding human genetics research and spoke out against its misuse and misappropriation, our findings make the case that scholars in the fields that have been shaped by Lewontin 1972 bear a moral and ethical responsibility to do the same.

## Acknowledgments

The authors wish to thank John Novembre, Doc Edge, Kevin Bird, and Emma Beyers-Carlson for their constructive comments on this manuscript. K.H. acknowledges support from NIH grant 1R35GM133428-01, a Searle Scholarship, a Sloan Research Fellowship, a Pew Biomedical Scholarship, and a Burroughs Wellcome Fund Career Award at the Scientific Interface.

